# Microbial functional guilds respond cohesively to rapidly fluctuating environments

**DOI:** 10.1101/2025.01.30.635766

**Authors:** Kyle Crocker, Abigail Skwara, Rathi Kannan, Arvind Murugan, Seppe Kuehn

## Abstract

Microbial communities experience environmental fluctuations across timescales from rapid changes in moisture, temperature, or light levels to long-term seasonal or climactic variations. Understanding how microbial populations respond to these changes is critical for predicting the impact of perturbations, interventions, and climate change on communities. Since communities typically harbor tens to hundreds of distinct taxa, the response of microbial abundances to perturbations is potentially complex. However, while taxonomic diversity is high, in many communities taxa can be grouped into functional guilds of strains with similar metabolic traits. These guilds effectively reduce the complexity of the system by providing a physiologically motivated coarse-graining. Here, using a combination of simulations, theory, and experiments, we show that the response of guilds to nutrient fluctuations depends on the timescale of those fluctuations. Rapid changes in nutrient levels drive cohesive, positively correlated abundance dynamics within guilds. For slower timescales of environmental variation, members within a guild begin to compete due to similar resource preferences, driving negative correlations in abundances between members of the same guild. Our results provide a route to understanding the relationship between functional guilds and community response to changing environments, as well as an experimental approach to discovering functional guilds via designed nutrient perturbations to communities.

## Introduction

Natural microbial communities, in contexts ranging from human hosts to soils, are buffeted by perturbations due to changes in host physiology, moisture, nutrients, pH, and temperature. Understanding how these environmental fluctuations impact community composition, interactions (*1*), and ultimately metabolic processes (*2*) is of critical importance. For example, changes in moisture, temperature, nutrients, and pH in soils impact the production of greenhouse gasses (*3–8*).

However, understanding the response of these consortia to environmental fluctuations remains a challenge because they harbor hundreds of distinct taxa. In principle, the abundance of each taxon might respond to perturbations via many distinct mechanisms. For example, changes in nutrient availability can impact interactions by relieving competition, while changes in moisture or pH can alter oxygen and nutrient availability respectively (*2, 9, 10*). Due to the diversity of taxa and mechanisms that can impact their abundances, one might expect that the response of a community environmental perturbations should be complex and high-dimensional.

Despite the potential complexity of community responses to perturbations, empirical and theoretical results suggest that communities are comprised of groups of taxa, or functional guilds, that perform similar functional roles. For example, in anaerobic digesters groups of taxa specialize in performing distinct steps in the fermentation cascade (*11, 12*). Similarly, in marine snow-degrading communities, groups of taxa perform polysaccharide degradation, oligomer uptake, and cross-feeding of excreted metabolites (*13*). These functional guilds are comprised of multiple coexisting taxa with similar metabolic preferences. As a result, while communities often retain hundreds of members, in many cases, these taxa can be grouped into functional guilds, with members of a guild exhibiting similar metabolic preferences. Functional guilds are thought to emerge, at least in part, from correlations in microbial traits such as strains specializing in sugar or acid catabolism (*14*) but not both.

Therefore, rather than dissecting how each strain responds to environmental perturbations, it might be much simpler to understand how guilds respond collectively to environmental perturbations (*15*). Since members of a single guild participate in similar metabolic transformations, it might be reasonable to assume guilds respond cohesively to environmental perturbations. Indeed, there is empirical evidence that functional guilds respond collectively to changing environmental conditions. For example, complex soil communities in bioreactors exhibit reproducible transitions between denitrification and dissimilatory reduction to ammonia as the carbon-to-nitrogen ratio is varied, reflecting changing dominant functional guilds in the system. This transition is thought to arise from the stoichiometric differences between denitrification and DNRA (*16*). Similar patterns are observed in large-scale metagenomic surveys of soil microbiomes (*1*). Likewise, distinct functional guilds in the cow rumen microbiome change in abundance in response to changes in lactate production during fiber fermentation (*17*). Theoretically, the notion of functional guilds has been described using a modular structure in the traits that members of a community possess (*18*).

Therefore, functional guilds potentially provide a route to understanding how communities collectively respond to environmental change. Specifically, rather than dissect the response of each strain in a system to a perturbation, it might be sufficient to understand how functional guilds respond to environmental changes (*19, 20*). In this sense, functional guilds might enable a more coarse-grained view of the response of communities to environmental change (*21*).

Here we investigate this idea using simulations and experiments. Counterintuitively, we find that the response of functional guilds depends on the timescale of environmental changes. Rapid changes in nutrient levels drive correlated dynamics between guild members, with members of the same guild increasing or decreasing their abundances cohesively across the guild. In this fast fluctuation regime, the community-level response reflects the guild structure. In contrast, when environmental fluctuations occur on a slow timescale, abundance dynamics are dominated by intra-guild competition. In this regime, the abundance of members of the same guild exhibit negatively correlated dynamics due to competitive interactions, and the community-level response does not reflect the guild structure.

## Results

We analyze a simple mathematical model of communities with functional guilds. Studying this model *in silico* and analytically reveals the primary finding of our study, that the timescale of environmental fluctuations determines whether or not members of a guild respond cohesively to these fluctuations.

### Trait-based description of microbial community

Here we study the response of communities to environmental change using a consumer-resource modeling (CRM) framework. The consumer-resource framework is a natural one for interrogating environmental changes because it directly couples the environment (resources) and growth. In the CRM framework, environmental fluctuations can be instantiated through time-varying resources.

A CRM framework naturally permits us to specify functional guilds by defining groups of taxa with similar metabolic resource preferences. Fig. 1A shows a schematic of a four-member community with two functional guilds (orange and blue) with strains that exclusively consume non-overlapping nutrients (shapes). Note that the orange strains compete for shared resources, but do not consume the resources utilized by the blue strains (and the converse).

**Figure 1:**
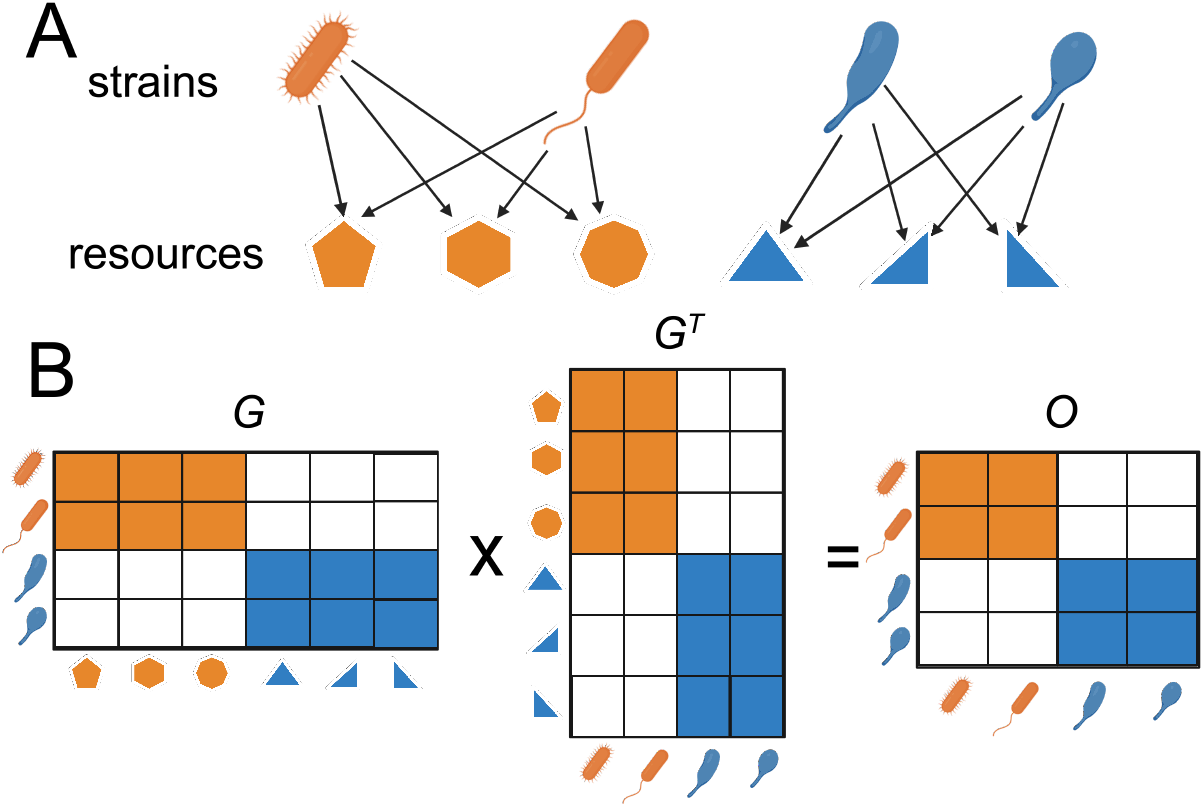
A model for resource consumption preferences and overlap. **(A)** We study a consumer-resource model, which consists of microbial strains (cartoon microbes) and the resources consumed (shapes). We consider a simple community with two functional guilds, represented by the colors. **(B)** The community description illustrated in (A) is formalized by a “growth matrix” *G*. The rows *i* of this matrix represent strains, the columns *α* represent resources, and the entries are the products of uptake rates and yields *r*_*i*,*α*_*1*_*i*,*α*_ (equivalent to the exponential growth rate in the high resource limit *R*_*α*_ *≫ R*_0_, Eq. 1). **(C)** The similarity of two strains can be quantified by their “overlap matrix” *O* = *GG*^*T*^.

The schematic in Fig. 1A can be formalized by defining a growth matrix, *G*, which specifies the growth rates of each strain on each resource (Fig. 1B). In particular, the rows of this matrix correspond to the growth rates of each strain on each resource 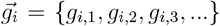, where the second index indicates the resource.

This formalism gives rise to a natural measure of metabolic similarity: the overlap matrix, *O* = *GG*^*T*^ (Fig. 1A,B). Each element in this matrix *O*_*i*,*j*_ corresponds to the dot product of the growth rate vectors of strains *i* and 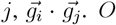 therefore quantifies the pairwise resource preference overlap of strains in the community, and its structure encodes the guild structure within the community. Groups of strains with high overlap (orange, blue blocks Fig. 1C) correspond to functional guilds of strains with similar resource preferences.

The resource preference matrix (*G*), resource supply rates, and death rates define the dynamics in our consumer-resource model as follows

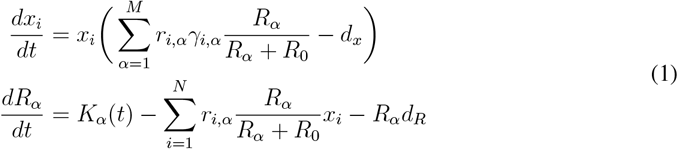

for *N* strains and *M* resources, where *x*_*i*_ are the consumers, *R*_*α*_ are the resources, *1*_*i*,*α*_ are the yields, *R*_0_ is the resource affinity, *d*_*x*_ is the consumer death rate, *K*_*α*_(*t*) are the time-dependent resource influx rates, and *d*_*R*_ is the resource depletion rate. Following (*22*), we have decomposed the growth rates *g*_*i*,*α*_ into the product of a resource uptake rate, *r*_*i*,*α*_, and biomass yield *1*_*i*,*α*_. Note that *r*_*i*,*α*_ has units of resource concentration per unit biomass per unit time and *1*_*i*,*α*_ has units of biomass per unit resource concentration, so *g*_*i*,*α*_ = *r*_*i*,*α*_*1*_*i*,*α*_ is a rate.

Crucially, in our model, the supply rates of the resources are time-dependent (*K*_*α*_(*t*)) whereas prior studies treat the supply rate of resources as time-independent and study the steady state of the community at long times (*23*).

### Abundance dynamics across timescales of environmental fluctuations

We simulate the abundance dynamics of a community subject to a range of environmental fluctuation frequencies. We randomly generate a community that has two functional guilds (Methods). Fig. 2A shows the *G* resource preference matrix used for these simulations. The community includes 20 strains consuming 100 different resources. The two functional guilds each comprise 10 taxa indicated by the orange and blue blocks in Fig. 2A. Unlike the example in Fig. 1, noise is added to this matrix, giving rise to members of the orange guild consuming some nutrients that are predominantly utilized by the blue guild and the converse. The role of this noise is explored below. Finally, the *G* matrix includes a set of private resources, one per strain, that ensures coexistence without biasing the composition within functional guilds (diagonal block, Fig. 2A). Fig. 2B shows the overlap matrix *O*, where the guilds become clear.

**Figure 2:**
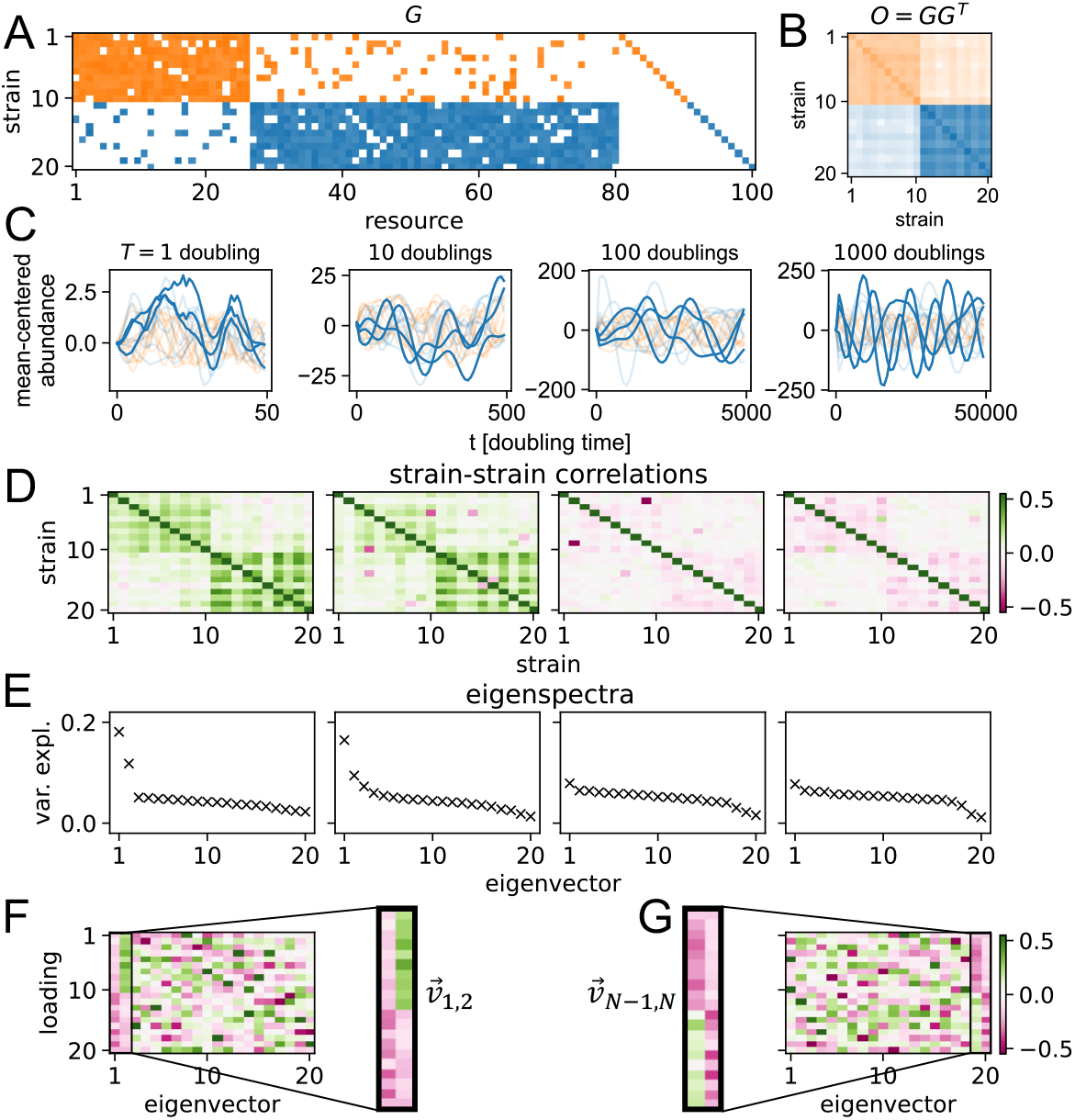
Microbial guilds interacting via resource competition are correlated on short timescales and anti-correlated on long timescales. **(A)** Matrix (*G*, Fig. 1) enumerating growth rates for strains (rows) on resources (columns). White entries indicate no growth. Colors indicate growth, with orange and blue demarcating the two guilds and the colormap varying linearly from 0 to max(*G*). *G* is corrupted by giving *p*_*f*_ = 0.1 probability for each entry of *G* to switch between 0 and *U* (⟨ *r*⟩ γ − Δ*G*, ⟨ *r ⟩ γ* + *1* + f:. *ΔG*), where *U* (*a, b*) denotes a uniform distribution on the interval (*a, b*). *r*, *1*, and f:.*G* are set to default values (Methods). Finally, each strain is given a private resource to ensure coexistence. **(B)** Overlap matrix *O* = *GG*^*T*^ quantifies the pairwise similarity of strain-strain resource preferences. Colors indicate guilds, and the colormap varies linearly from 0 to max(*O*). **(C)** Simulated mean-centered abundance dynamics are shown across four timescales. The title gives the timescale *T* = 2π/ ⟨ *ω*⟩ of environmental fluctuations in units of average doubling time on a single resource, where ⟨ *ω*⟩ is the average resource fluctuation rate. Abundance dynamics are colored according to guild and dynamics from strains 11, 12, and 17 are highlighted to illustrate the loss of intra-guild cohesion as the timescale increases. Note that axes scales are changed in each panel so that dynamics are visible. **(D)** Pearson’s correlation coefficient for the abundance dynamics of each strain pair is shown at each fluctuation timescale (Methods). **(E)** Rank-ordered variance explained by the eigen-vectors of the corresponding correlation matrix in (D). **(F)** Eigenvectors for *T* = 1 doubling case (left panels, (D),(E)), ordered according to the rank of the eigenvalues. The eigenvectors corresponding to the two largest eigenvalues reflect the block structure. In particular, the strains in the blue block are weighted most heavily in 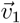, and the strains in the orange block are weighted most heavily in 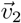. The corresponding eigenvalues are above the background (E, left panel), indicating a cohesive intra-guild response to fast environmental fluctuations. **(G)** Eigenvectors for *T* = 1000 doublings, ordered according to the rank of the eigenvalues. The eigenvectors of the two smallest eigenvalues reflect the block structure. In particular, the strains in the blue block are weighted most heavily in 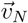, and the strains in the orange block are weighted most heavily in 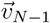. These are the smallest eigenvalues, indicating a there is no cohesive intra-guild response to environmental fluctuation for slow fluctuations.

The dynamics of the community are governed by Eq. 1, with environmental fluctuations taking the form of temporal changes in the input resource vector 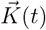. To introduce environmental fluctuations at different timescales, the input rate of each resource *K*_*α*_ fluctuates as a sinusoid.

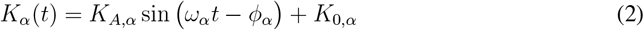

For a given simulation, we set a timescale of fluctuations via the frequencies *ω*_*α*_ which vary by 10% across resources around the average value (Methods). The average resource fluctuation frequency (⟨*ω* ⟩) defines the timescale of environmental fluctuations, *T* = 2π/ ⟨*ω* ⟩.

Such periodic fluctuations are qualitatively similar to many environmental dynamics experienced by natural communities, such as diel cycles (*24*), tidal patterns (*25*), and seasonal variation (*26*). Since each resource fluctuates at a slightly different frequency, these fluctuations result in a continuously changing environment with a characteristic timescale set by ⟨*ω* ⟩. To study the response of the system at different timescales, we vary this average frequency.

Abundance dynamics (colored by guild identity) were simulated for environmental fluctuations across a range of timescales (Fig. 2C). We define timescales in terms of the average doubling time across all members of the community. Fast fluctuations have a timescale *T* which is on the order of the average doubling time in the community (Fig. 1C, left two panels). Slow environmental fluctuations mean that *T* is much longer than the average doubling time (Fig. 2C, right two panels). Examination of Fig. 2C suggests that at fast timescales (left panels) of environmental fluctuations, members of the same guild exhibit cohesive dynamics, with abundances of taxa within the same guild rising and falling together in response to environmental changes. In contrast, when environmental fluctuations are slow, the abundances of members of the same guild do not exhibit cohesive dynamics (Fig. 2C, right panels).

### Functional guilds dominate response for fast but not slow environmental fluctuations

To formalize the qualitative observations in Fig. 2C, we computed correlations of abundances across each pair of strains in the community (Fig. 2D; Methods). For fast environmental fluctuations, we observe positively correlated abundance dynamics of members of the same guild (green blocks, Fig. 2D, left panels) and converse for long-timescale fluctuations (pink blocks, Fig. 2D, right panels).

### Fast environmental fluctuations drive correlated, cohesive response

When environmental fluctuations are fast, the community does not have time to approach a steady state. In this case, the strain abundance dynamics are dominated by the environment defined by the resource influx rates 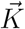. Strain abundances respond to changes in the availability of their preferred resources. This leads to the correlations in abundance dynamics of similar strains such as those shown in Fig. 2C and D. This cohesive response is also reflected in the spectrum of the abundance correlation matrix, 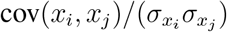, in the quickly fluctuating regime (Fig. 2E, left two panels). The first eigenvalue captures the dynamics of the large (blue) guild, whereas the second eigenvalue captures the dynamics of the smaller (orange) guild. This can be seen in the eigenvectors (Fig. 2F), where the first eigenvector has high magnitude loadings for strains in the blue guild, whereas the second eigenvector has high magnitude loadings for strains in the orange guild.

Overall, the response of the community to fast environmental fluctuations matches the structure of the trait matrix, leading to a cohesive response at the guild level.

### Slow environmental fluctuations cause competition to drive non-cohesive response

The environment-dominated regime can be contrasted to the regime defined in the opposite limit – very slow environmental fluctuations. In this regime, the strain abundances equilibrate much more quickly than the resource environment changes, so the system is always near steady state. Therefore, the competitive interactions between the strains dominate the abundance dynamics. This leads to negative correlations between members of the same guild because they consume very similar resources (Fig. 2D, right panels). This can also be seen in the eigenspectrum of the correlation matrix and corresponding eigenvectors (Fig. 2E, right panels), which is the inverse of the eigenspectrum and modes in the fast fluctuation limit. Figs. 2E and G show that the modes encoding the guild structure correspond to the two smallest eigenvalues. This decrease in variance explained by the block structure relative to the rest of the eigenspectrum shows that the response is not cohesive and that the growth of strains within a guild inhibits the growth of other strains within that guild.

### Analytic calculation connects covariance to guild structure

Finally, we sought to formalize the connection between guild structure and the strain-strain correlation matrix. We show analytically that a scaled version of the overlap matrix *O* = *GG*^*T*^ provides an approximation to the expected covariance between strains in the community (Supplementary Information). This is consistent with our empirical results (Fig. 2) and reveals the mathematical basis for the observed relationship between guild structure and strain-strain abundance correlation dynamics.

### Cohesion of guilds is driven by the strength of guild structure and rate of environmental fluctuations

Next, we asked what factors control the cohesion of the functional guilds in response to environmental fluctuations. In addition to the timescales of environmental fluctuation noted above, we hypothesized that the cohesive response requires the well-defined block structure of the trait matrix *G*.

To examine this effect, we simulate the system described in Eq. 1 while varying two parameters: the flip probability *p*_*f*_, which is the probability that each growth rate *g*_*i*,*α*_ deviates from the block structure (quantifying the level of guild structure of the trait matrix *G*), and the fluctuation timescale *T* = 2*π*/⟨*ω* ⟩. Thus increasing *p*_*f*_ corresponds to a weakening of the guild structure. Fig. 3 shows the results, where the cohesion of the community response is quantified by calculating the average intra-guild strain-strain correlations ⟨ *ρ*_*g*_ ⟩ averaged across 10 simulated correlation matrices and randomly generated environments (Methods). On fast timescales (*T* = 1, 10), we see a monotonic decrease in the average correlation as *p*_*f*_ increases. We conclude that the cohesion of guilds in response to environmental perturbations is modulated by the strength of the guild structure, with stronger guilds resulting in more cohesive responses at fast timescales.

**Figure 3:**
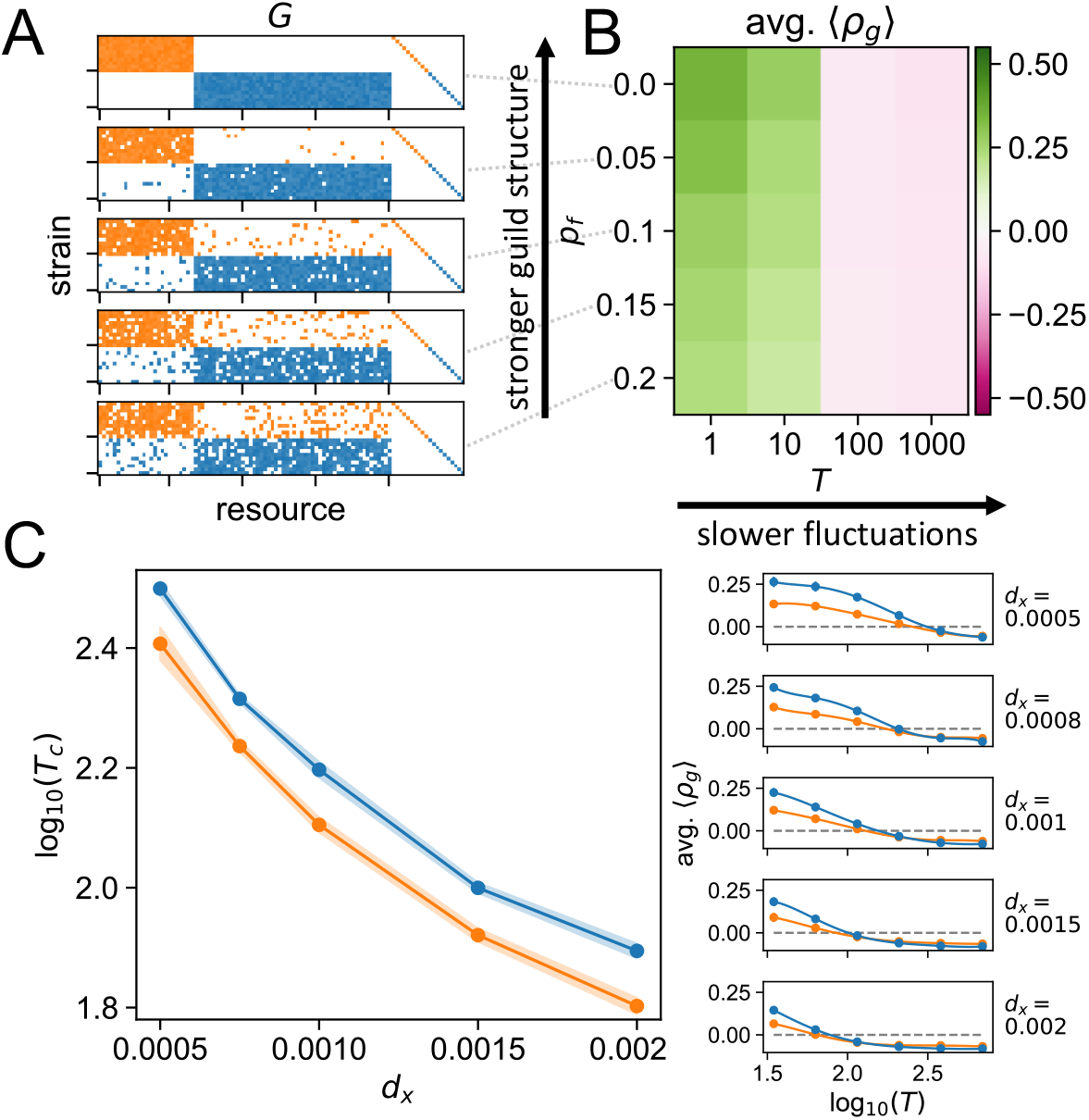
Intra-guild cohesion is driven by resource utilization structure and perturbation timescale. **(A)** Examples of growth rate matrices *G* corresponding to different values of *p*_*f*_ (dashed lines). **(B)** Average intra-guild strain-strain correlations ⟨ *ρ*_*g*_ ⟩in simulated abundance correlation matrix (for the blue guild) averaged across 10 simulations for different values of *p*_*f*_ and environmental fluctuation timescale *T* (Methods). Higher average ⟨ *ρ*_*g*_ ⟩ indicates a higher level of cohesion within the guild. **(C)** Death rate sets the timescale of crossover from positive to negative intraguild correlations. The left panel shows the logarithm of the crossing timescale log(*T*_*c*_), plotted as a function of death rate *d*_*x*_. Shaded region indicates uncertainty (Methods). *T*_*c*_, which can be interpreted as the characteristic timescale of the system, decreases as *d*_*x*_ increases. Average growth rate ⟨*g*⟩ is set to 1 by changing average uptake rate ⟨*r⟩* to 5 and keeping yields *γ*_*i*,*α*_ at the default value of 0.2, so death rate can be interpreted as a fraction of the growth rate. All other parameter values are set to the default values (Methods). The right panels show the average ⟨ *ρ*_*g*_ ⟩as a function of the logarithm of the environmental fluctuation timescale log(*T*) for different death rates *d*_*x*_. Errorbars indicate standard deviations across 10 simulations.

### Death rate sets the timescale of the system

Finally, we asked what biological parameters set the timescale of the ecological processes. In particular, we defined this timescale as the environmental fluctuation period at which average ⟨ *ρ* _g_⟩ switched sign, *T* = *T*_*c*_. We then repeated our simulations with *p*_*f*_ = 0.1 for different values of the growth rate, death rate, and intra-guild variance in growth rate. The intra-guild variance in growth rate was defined as the width of the distribution from which non-zero growth rate values were drawn (Methods). We then fit the average intra-guild correlation as a function of the fluctuation period to a smoothing spline to infer *T*_*c*_ for each parameter set (Fig. 3C, right panels; Methods).

We find that the death rate sets *T*_*c*_. Both guilds show decreasing *T*_*c*_ as the death rate increases (Fig. 3C). In our simulations, *T*_*c*_ does not depend on the growth rate or variance in intra-guild growth rate (Fig. S1), although this empirical treatment cannot rule out this possibility in different parameter regimes. This result indicates that the transition from a cohesive, environmentally-dominated regime to a non-cohesive, competition-dominated regime is set by how quickly organisms die. Ecologically, this is intuitive: the death rate effectively sets the turnover rate of the community and therefore the timescale on which ecological processes occur (*27*).

### Experimental demonstration of dynamic dependence of guild cohesion

Next, we wanted to test the proposal that intra-guild abundance dynamics are cohesive on short timescales and not cohesive on longer timescales. Experimentally, imparting sinusoidal resource fluctuations is a challenge. However, performing serial batch culture experiments, where communities are grown for fixed intervals before being diluted into fresh nutrients, is routine (*28*).

Therefore, we set out to design a serial batch culture experiment, with synthetic communities, where the *G* matrix is inferred from experimental data and the environment can be manipulated. To do this, we assayed the growth phenotypes of ~20 strains on 10 distinct carbon sources (*29*) in monoculture. Second, we constructed a synthetic community from these isolates, and rather than fluctuating resources over time, we varied the resources present in the community. We repeated this experiment for many combinations of resources. Below we show how the dynamics of this experiment exhibit qualitatively similar behavior to the situation analyzed above.

### Timescale dependence of cohesion across environments

To relate our theoretical result to an experimental batch serial-dilution context, we redefine timescale of environmental fluctuations as the length of time that has passed following an environmental change. We therefore start our two-guild communities at equal abundances, subject them to many randomly generated environments, as in the experiment illustrated in Fig. 4A, and calculate correlations between strain abundances across environments as a function of time. To mimic experimental serial-dilution conditions, we supply nutrients to the community in periodic batches and dilute after a fixed time. The model then becomes

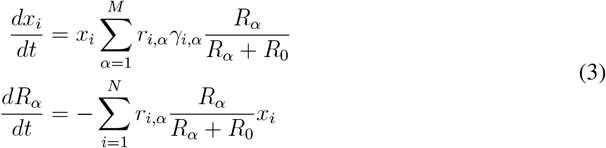

with initial conditions defined for each batch cycle by

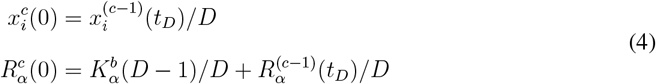

where *c* indexes the cycle, *D* is the dilution factor during serial dilution (taking the place of death rates *d*_*x*_ and *d*_*R*_ in Eq. 1), mimicking a dilution constant in batch culture experiments (*1*). 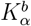 is the concentration of resource *α* supplied in the fresh media at the beginning of each cycle. 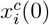 and 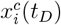 are the values at the beginning and end of cycle *c*. 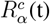 is defined in the same way.

**Figure 4:**
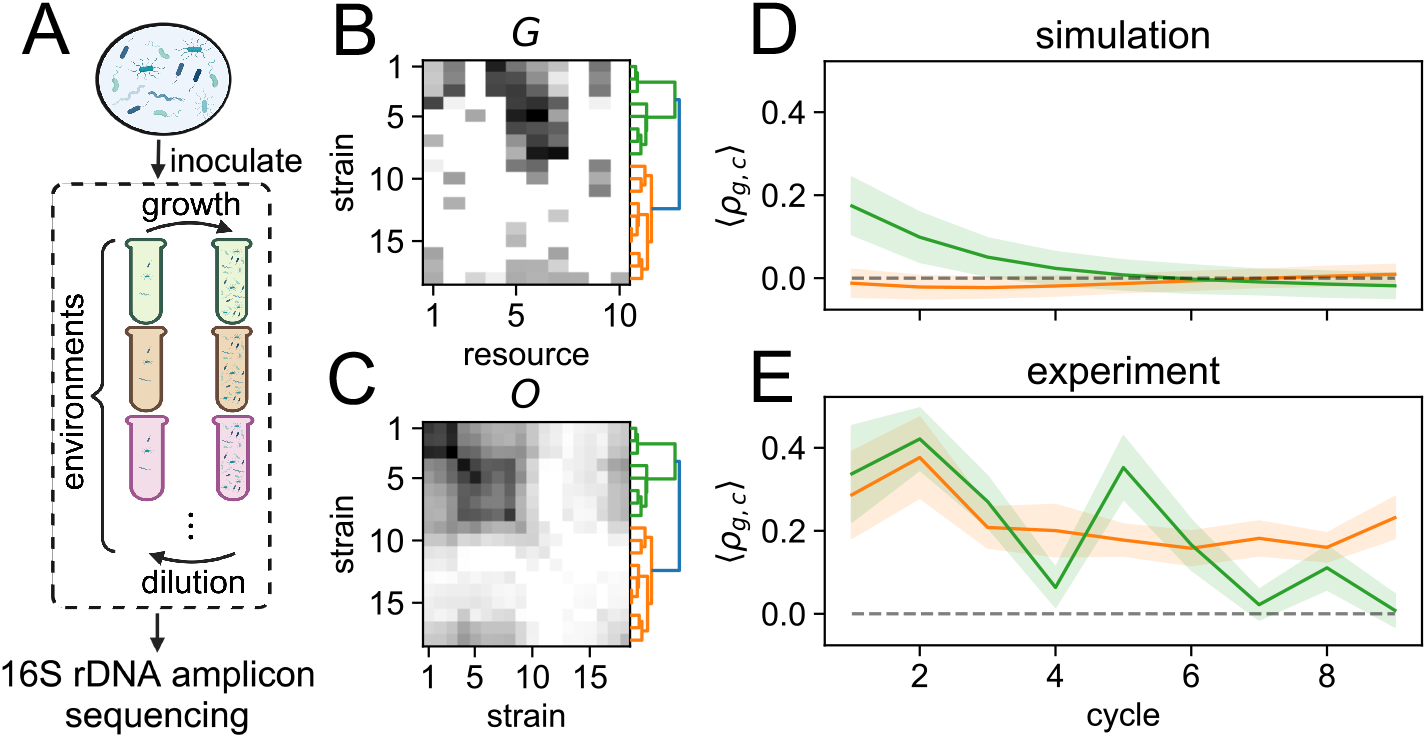
Experimental demonstration of decreasing guild cohesion over time during serial batch culture. **(A)** Experimental schematic. A synthetic community with 18 strains was inoculated into 32 environments, each comprised of a number of randomly chosen carbon resources. This community was passaged for 9 batch growth cycles, with a 10-fold dilution into fresh media at the end of each cycle. **(B)** Prior to the experiment, the *G* matrix was estimated by growing each strain independently on each of the 10 carbon sources (Methods). Entries are growth rates measured via OD in time in a plate reader, with the colormap varying linearly from 0 to max(*G*) = 0.7 h^*-*1^. Hierarchical clustering is used to identify guilds (green and orange branches in dendrogram). **(C)** Inferred overlap matrix *O* of the synthetic community. Guilds are shown by the dendrogram, and the colormap is varied linearly from 0 to max(*O*)= 1.1 h^*-*2^. **(D, E)** Time-series of the simulated (panel D) and experimental (panel E, measured by sequencing, Methods) average intra-guild correlation for the synthetic community in the experimental environments. Variance is calculated by resampling with replacement and shown by the shaded region (Methods). Green and orange correspond to the green and orange guilds, respectively (panels B and C). In the simulation, parameters *D* and 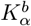 are adjusted from default values to match experimental values (Methods).

We numerically integrate this model for randomly generated versions of the two guild communities used above (Fig. S2A,B; Methods). Next, we calculate strain-strain abundance correlations across an ensemble of 100 randomly generated environments as a function of serial dilution cycle (Methods). Consistent with the continuous case examined previously, average intra-guild correlations decrease as a function of time, here defined as the cycle at which correlation was measured (Fig. S2C).

### Resource competition interactions result in correlations at short times and anti-correlations at long times *in vitro*

To test the implications of our theoretical results, we conducted an experiment to measure correlations between strains competing for resources across time. We grew a synthetic community consisting of 20 strains with known carbon utilization capabilities (*29*) for 9 cycles in serial batch culture in 32 different, randomly generated environments. Next, we used 16S sequencing with a spike-in to measure the absolute abundances of each strain at the end of each cycle (Methods).

To compare the experiment to the *in silico* expectation, we first need to define experimental guilds. For the 18 strains with measured abundances in all cycles, we used hierarchical clustering to identify two functional guilds (Fig. 4A,B; Methods). Next, we integrate Eq. with the resources provided in the experiment and the inferred *G* matrix. Finally, we plot the average intra-guild correlation ⟨ *ρ*_*g*,*c*_⟩ for each inferred guild *g* at each cycle *c* in both the simulation and experiment.

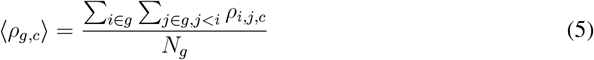

where *ρ*_*i*,*j*,*c*_ is the Pearson’s correlation coefficient across environments between strains *i* and *j* at cycle *c* (Methods), and *N*_*g*_ is the number of strain pairs in the guild.

For the guild that consists of strains that can utilize many of the carbon resources (Fig. 4D,E; green curves), we observe the characteristic decrease in ⟨ *ρ*_*g*,*c*_⟩ associated with functional guilds interacting via resource competition. This is consistent with our theoretical expectation (Fig. S2C). Crucially, this recapitulates the basic intuition from Fig. 2, that strains within a guild are positively correlated for short timescales of environmental variation and the converse.

For the other guild (Fig. 4D,E; orange curves), we do not see a significant decrease in ⟨ *ρ*_*g*,*c*_⟩ across cycles in either the simulation or experiment. Examination of the resource utilization of the strains in this guild, however, shows that they do not grow quickly on *any* resources relative to the strains in the green guilds, and that the resources they do consume are also consumed by these guilds (Fig. 4B). This suggests that these strains are unlikely to be able to survive by competing for resources. While this is the case *in silico* (Fig. S3A), strains in the orange guild are able to survive *in vitro* (Fig. S3B). To account for this discrepancy between experiment and simulation, we note that the simulation assumes that resource competition is the only mechanism of interaction between strains in the community. We speculate, therefore, that strains in the orange guild survive due to a cross-feeding interactions with strains in the green guild, perhaps mediated by overflow metabolism (*30*). This is supported by persistent positive correlations between strains 2 and 3 in the green guild and many of the strains in the orange guild (Fig. S4). This is consistent with theoretical expectations for strains that interact via cross-feeding (Supplementary Information; Fig. S5).

## Discussion

Here, we have characterized the response of microbial communities composed of metabolic guilds to environmental fluctuations. We demonstrate that fast environmental fluctuations excite a cohesive response within metabolic guilds and that cohesion is lost for slow environmental fluctuations. The changing cohesion of metabolic guilds with timescales of fluctuations has important implications for understanding community dynamics in the wild. For example, it is routine to study correlations in abundances of microbes across time and space (*31*). Interpreting correlated abundance dynamics is a long-term challenge in microbial ecology (*32, 33*). Here we show that the sign of these correlations can vary strongly with the timescale of environmental variation driving abundance dynamics. This means that care must be used in interpreting when abundance correlations might reflect ecological associations (*34*). The observation also offers an opportunity to understand metabolic guild structure in communities and to expose underlying structural properties that drive community function.

### Impact of timescale on collective properties

This result illustrates the importance of dynamics in the collective properties of microbial communities. In this work, we define timescale in two different ways: (1) the rate of environmental fluctuation (Fig. 2, 3) and (2) time after environmental perturbation (Fig. S2,Fig. 4). In both cases, the community response is the same, cohesive on short timescales, and not cohesive on long timescales. In this context, we should expect that on short timescales the collective properties of a community (e.g. nutrient flux) should contain contributions from each organism within a functional guild. On long timescales, however, the collective properties may be dominated by a single strain that is the most effective competitor within the guild in that environment. As a result, we might expect that the collective flux of metabolites (e.g. (*2*)) might reflect the activity of all members in a guild on short timescales and be driven by the dominant competitor within the guild on longer timescales. It remains open to test this hypothesis in microcosm experiments with controlled environmental forcing (*28, 35–37*).

### Implications for experimental determination of functional guilds

In addition to providing insight into natural ecological processes, this work informs experimental design. In particular, the inference of functional guilds has recently been a topic of interest in microbial ecology and microbiology (*38–40*). Identification of functional guilds in microbial communities is valuable because it can enable coarse-graining communities using effective groups of taxa (*2, 41, 42*). This coarse-graining provides a low-dimensional description of communities that can enable predictions of community dynamics and function.

A common approach to inferring guilds is to perform serial dilution enrichments to identify strains that perform well in an environmental condition of interest (*28*). Our theoretical results suggest that the observed correlations between strains in such an experiment depend on the timescale over which those correlations are observed. Thus, ideally, one would observe correlated responses of groups over short timescales. We proposed and tested an experimental procedure for identifying functional guilds via serial-dilution experiments. One can imagine this approach being scaled up by performing short-term enrichment experiments on complex communities in diverse environments (*43*). In these contexts, our approach might help define metabolically cohesive units within the community.

### Extension to cross-feeding interactions

Here, we largely restricted ourselves to consideration of resource competition. Recent work suggests that such interactions dominate in many contexts (*44, 45*). However, it is the case that metabolic byproducts from one organism can be used by another, and these cross-feeding interactions are qualitatively distinct from competition (*1, 17*). Intriguingly, we find that simulations with cross-feeding interactions give rise to positive *inter*-guild correlations for *all* fluctuation timescales (Supplmentary Information). This raises the possibility that different classes of interactions can be identified by studying correlations across timescales of environmental fluctuation. The role of cross-feeding in driving dynamics between guilds is an important avenue for future work.

### Interpretation and effect of private resources

In this study, we provide each strain with a fluctuating “private resource” that ensures that it maintains some significant biomass (Fig. 1A). This ensures that the community remains relatively stable over time, which is generally the case in naturally-occuring communities (*31, 46–48*).

Ecologically, such private resources need not be interpreted literally as nutrients available only to a single guild member of the community. Instead, these resources may be viewed more abstractly as some mechanism that is orthogonal to the block resources that maintains diversity within functional guilds, such as a spatial structure or phage predation (*49–52*). For instance, the background fluctuations introduced by the private resources may arise from migration from different environments or adherence to surfaces that provide unique access to resources (*53, 54*).

Although it is in principle possible for block resources alone to support coexistence, identifying a community that coexists across an ensemble of fluctuating resources requires significant finetuning. In particular, we expect that even *in silico* communities chosen such that members coexist at average values of the block resources will experience extinctions in fluctuating environments. Although it is in principle possible that such fine-tuning will arise from ecological and evolutionary processes, it necessarily biases the community composition and implicitly makes an assumption about the source of community stability. For this reason, we chose to maintain coexistence using private resources, allowing us to choose an unbiased distribution of traits within functional guilds. While extensive theoretical progress has been made regarding the source of community stability and strain coexistence (*55–57*), the effect of such processes on community cohesion dynamics has not been considered. Our results suggest that this is an exciting avenue of future research, raising questions about how assumptions about mechanisms of coexistence influence the dynamics of intra-guild cohesion.

## Acknowledgments

We thank Madhav Mani, Neelima Sharma, and members of the Kuehn lab for useful discussions.

## Author contributions

K.C., A.M., and S.K. conceptualized the research. K.C. performed the simulations. A.S. performed the analytic calculations and provided feedback on simulation results. R.K., K.C., and S.K. designed the experiments. R.K. performed the experiments. K.C., S.K., and A.S. wrote the manuscript with contributions from R.K..

## Data and materials availability

Code and data associated with this manuscript, including code and data sufficient to reproduce all figures, are publicly available at 17605/OSF.IO/J8S2V.

