## Supplementary material for "Microbial functional guilds respond cohesively to rapidly fluctuating environments": Methods and Supplementary Material

#### Consumer-resource model simulations

Consumer-resource model simulations were performed via the numerical integration of Equations 1, 4, and 18 using the implicit Runge-Kutta method "Radau" implemented in SciPy (*58*).

##### Random generation of environmental fluctuation rate and phase

To generate environmental fluctuations, we parameterize the resource influx rates

$$K_\alpha(t) = K_{A,\alpha} \sin\left(\omega_\alpha t - \phi_\alpha\right) + K_{0,\alpha} \tag{6}$$

by randomly drawing $\omega_\alpha$ from a uniform distribution from from $\langle\omega\rangle - 0.1\langle\omega\rangle$ to $\langle\omega\rangle + 0.1\langle\omega\rangle$, where $\langle\omega\rangle$ is the average fluctuation frequency. The phases $\phi_\alpha$ are drawn from a uniform distribution from $-\pi$ to $\pi$.

##### Random generation of $G$

The growth rate matrix $G$ was generated by first choosing average uptake rates $\langle r\rangle$ and a constant yield $\gamma$. Uptake rate and yield matrices were then constructed such that

1. Each element had a probability of $p_f$ to deviate from the block structure (non-zero uptake rate and yield outside of the blocks, zero uptake rate and yield inside of the blocks).
   (a) By default, $p_f = 0.1$, but this value is changed as indicated for the simulations associated with 3A and B.
2. Each non-zero uptake rate is drawn from a uniform distribution from $\langle r\rangle - \Delta G/\gamma$ to $\langle r\rangle + \Delta G/\gamma$, so that the growth rate varies from $\langle g\rangle - \Delta G$ to $\langle g\rangle + \Delta G$.
   (a) By default, $\Delta G = 0.1\langle r\rangle\gamma$, but this value is changed as indicated for the simulations associated with Figs. 3 and S1.

#### Random generation of batch environments

To randomly generate environments for the batch culture simulations shown in Fig. S2, each initial resource value $K_{B,\alpha}$ was set to either $0$ or the default value of $1$, each with probability $0.5$. Note that this differs from the choice of experimental environments and the corresponding simulation of the experiment (Figs. 4 and Fig. S3). These are described in the experimental methods section below.

#### Calculation of strain-strain correlation matrix across time

To calculate the strain-strain correlation matrices in continuous simulations (Figs. 2 and 3), the following procedure was used. First, strain abundances were initialized at their steady state values corresponding to constant resource influx at the average values $K_\alpha(t) = K_{0,\alpha}$. This was done to eliminate transient abundance dynamics at the beginning of the simulation. Next, abundance values for each strain were calculated for the fluctuating environment described in Eq. 6 with timesteps equal to the environmental fluctuation timescale $T$ ($\Delta t = T$). Finally, an array consisting of $1000$ timepoints was passed to the NumPy function `corrcoef` (*59*) to compute the correlation matrix.

#### Calculation of average $\langle \rho_g \rangle$

To calculate average $\langle \rho_g \rangle$ in Fig. 3, the strain-strain correlation matrix $\rho_{i,j} = \text{cov}(x_i, x_j)/(\sigma_{x_i}\sigma_{x_j})$ over time was calculated (see the previous section) for $10$ randomly generated $G$ matrices with randomly generated fluctuation rates $\omega_\alpha$ for each set of parameters. For each simulation, the average correlation was calculated across all pairs of strains ($N_g$ total pairs) in each guild. The average of these pairs $\langle \rho_g \rangle = \frac{\sum_{i\in g}\sum_{j\in g, j<i}\rho_{i,j}}{N_g}$ was then averaged across simulations (instantiations of $G$ and $\omega_\alpha$) to compute average $\langle \rho_g \rangle$.

#### Calculation of $T_c$

To determine the fluctuation rate at which average $\langle \rho_g \rangle$ switched sign, $T_c$, we fit average $\langle \rho_g \rangle$ to a smoothing spline with degree $k = 5$ (*60*). We then used the spline fit to interpolate between simulated values of average $\langle \rho_g \rangle$. $T_c$ was then defined as the average period at which the interpolated average $\langle \rho_g \rangle$ was $0$. Uncertainty in $T_c$ was calculated by resampling simulations with replacement at each fluctuation rate $1000$ times, recalculating $T_c$, and calculating the standard deviation of this ensemble.

##### Simulation parameter values

Unless otherwise specified, the parameter values for the simulations are

Table 1: Default simulation parameter values

| Parameter | Continuous Simulation (Eq. 1) | Batch Simulation (Eq. 4) |
|---|---|---|
| $\langle r \rangle$ | 1 | 1 |
| $\gamma_{i,\alpha}$ | 0.2 | 0.2 |
| $R_0$ | 0.01 | 0.01 |
| $d_x$ | 0.001 | NA |
| $d_R$ | 0.02 | NA |
| $K_{A,\alpha}$ | 10 | NA |
| $K_{0,\alpha}$ | 100 | NA |
| $K^b_\alpha$ | NA | 1 |
| $D$ | NA | 8 |
| $\Delta G$ | $0.1\langle r \rangle \gamma$ | $0.1\langle r \rangle \gamma$ |
| $p_f$ | $0.1$ | $0.1$ |

#### Experimental methods

##### Characterization of bacterial growth

We characterized growth of $20$ bacterial strains by growth in minimal media supplemented with different carbon resources. Each strain was revived from a glycerol stock and grown in $96$-deep well plates in $1.2\,\mathrm{mL}$ of $0.2$X Tryptic Soy Broth (TSB) at $30$°C with shaking for $2$ days and was subsequently back-diluted into $0.2$X TSB for $1$ more day. Cultures were then washed in carbon-free minimal media and used to inoculate $700\,\mathrm{uL}$ of a defined minimal media supplemented with different carbon resources at a low optical density ($OD_{600} = 0.01$) in $48$-well optical plates. The minimal media consists of $15\,\mathrm{mM}$ ammonium as the assimilatory nitrogen source, $40\,\mathrm{mM}$ phosphate buffer with the final medium pH adjusted to $7.3$, trace metals and vitamins, and $25\,\mathrm{mM}$ of carbon atoms from one of $10$ carbon compounds: arabinose, butyrate, deoxyribose, glucuronic acid, glycerol, mannitol, mannose, melibiose, propionate, and raffinose. Minimal media cultures were grown at $30$°C with shaking ($500\,\mathrm{rpm}$, Fisherbrand™ Benchtop Incubating Microplate Shaker) for $72$ hours with logarithmic sampling to measure optical density ($OD_{600}$). Growth rates $g_{i,\alpha}$ were inferred via a linear fit to $\log(OD_{600})$ in exponential phase. Biomass yield per carbon atom was inferred from endpoint $OD_{600}$ and known input carbon concentration ($\gamma = \Delta OD/\Delta C$, where $\Delta OD$ is the change in $OD_{600}$ during growth cycle and $\Delta C$ is the carbon concentration consumed assuming complete consumption). Finally, uptake rate was calculated using the inferred $g_{i,\alpha}$ and $\gamma_{i,\alpha}$ as follows: $r_{i,\alpha} = g_{i,\alpha}/\gamma_{i,\alpha}$. We note that this calculation is approximate and may be corrupted by incomplete consumption of the carbon source. Growth rates and yields for all strains used for the experiment can be accessed in the code and data repository associated with this manuscript, https://doi.org/10.17605/OSF.IO/J8S2V.

**Serial-dilution batch culture experiment**

The $20$ characterized bacterial strains were revived from glycerol stocks as described above and grown to early stationary phase before being used to inoculate $32$ media conditions with distinct carbon resource profiles. Each media condition was made with the minimal media described above, amended with a total of $25\,\mathrm{mM}$ carbon atoms from a unique combination of $1$ to $6$ carbon resources. Each media condition was inoculated with a uniform mixture of the $20$ strains at a starting total $OD_{600} = 0.01$ in $700\,\mathrm{uL}$ in a $48$-well optical plate sealed with a breathable film seal. These batch culture plates were grown at $30$°C with shaking ($500\,\mathrm{rpm}$, Fisherbrand™ Benchtop Incubating Microplate Shaker) for $48$ hours. At the end of $48$ hours, each batch culture was diluted into fresh media with a $1{:}10$ dilution. The remaining culture was spun down and frozen for downstream DNA extraction.

#### Choice of environments

32 media conditions, representative of 32 environments, were randomly generated as follows: given a total of 10 carbon resources, each resource had a 25% chance of being selected for a given environment, resulting in environments with different numbers of carbon resources. The total carbon atom concentration in every environment was fixed to $25\,\mathrm{mM}$ and evenly partitioned between the resources assigned to that environment. The robustness of the community composition response to this environment selection protocol was assessed using 5 replicate consumer-resource model simulations of a batch culture instantiated with the 20 strains and measured $G$ used in the experiment (Eq. 4). All five replicate simulations demonstrated qualitatively similar responses in the community composition, so a representative set of environments was chosen for the experiment (in particular, the environments corresponding to the first simulation). Simulation results and the set of environments used for the experiment can be accessed in the code and data repository associated with this manuscript, https://doi.org/10.17605/OSF.IO/J8S2V.

#### Sequencing

Genomic DNA was extracted from bacterial cell pellets using the Qiagen DNeasy Blood and Tissue plate-based kit with a modified lysis protocol for Gram-positive bacteria recommended in the kit. To estimate the absolute abundance of bacterial 16S rDNA amplicons, we added known quantities of genomic DNA extracted from *Escherichia coli* K-12 (sourced from the Duchossois Family Institute Commensal Isolate Library, Chicago, IL, USA) to the pellet resuspension buffer prior to DNA extraction. 16S amplicon sequencing was performed using the Illumina 16S Metagenomic Sequencing Library Preparation protocol with some modifications. The V3-V4 region was amplified using forward primer 341-b-s-17 with Nextera adapter (TCGTCGGCAGCGTCAGATGTGTATAAGAGACAGCCTACGGGNGGCWGCAG) and reverse primer 806R_Apprill with Nextera adapter (GTCTCGTGGGCTCGGAGATGTGTATAAGAGACAGGGACTACNVGGGTWTCTAAT). Sequences were generated on the Illumina Miseq platform using a 2 x 300 bp paired-end v3 reagent kit with a 10% PhiX spike-in (Illumina, San Diego, CA, United States).

#### Sequence data processing and ASV-assignment with DADA2

The following pre-processing steps were performed in DADA2 (*61*). Raw Illumina reads were stripped of primer sequences, truncated after a Phred quality score below 5, and filtered to a maximum expected error of 2 in the forward reads and 5 in the reverse reads based on Phred scores. ASV assignment on processed reads was performed using DADA2 and the recommended analysis pipeline. Forward and reverse reads were denoised separately, then merged and filtered for chimeras. ASV inference was performed by pooling all samples together. After processing, each strain was assigned to one ASV (based on prior 16S rDNA assignment by Sanger Sequencing performed on axenic cultures of each isolate) and one strain was assigned to two ASVs that were correlated across samples. The sequencing depth was $10^3$-$10^4$ reads per sample. To obtain the absolute abundance of each ASV per sample, the ASV counts in each sample were divided by the spike-in *Escherichia coli* K-12 ASV counts in the same sample.

#### Identification of functional guilds

Functional guilds were identified by first creating an empirical growth rate matrix ($G$, strain x carbon resource) on a synthetic community, using measured growth rates. Next, this matrix was hierarchically clustered using Ward's method (*62*). Individual guilds were defined using a distance threshold on the resulting dendrogram (95% of the maximum link distance).

#### Calculation of strain-strain correlations across environments $\rho_{i,j,c}$

To calculate the strain-strain correlation matrices at each cycle in batch conditions (Figs. 4 and S2), the following procedure was used. First, abundance values for each strain were measured or calculated at the end of each cycle. Next, an array consisting of abundance values for each strain in each environment was passed to the NumPy function `corrcoef` (*59*) to compute the correlation matrix for each cycle.

**Uncertainty calculation of batch intra-guild correlation coefficient $\langle\rho_{g,c}\rangle$**

To calculate the uncertainty of $\langle\rho_{g,c}\rangle$ (where brackets indicate averaging over $i, j \in g$, Eq. 5) in Figs. 4D,E and S2C, environments are first resampled with replacement $1000$ times. $\langle\rho_{g,c}\rangle$ is calculated for each resampled set of environments, and the standard deviation of this ensemble is shown by the envelope.

### Supplementary Information

#### Analytic approximation of the consumer covariance matrix

To derive the expected covariance matrix of strain abundances in slow fluctuation limit, we first rewrote Eq. 1 in matrix form as follows.

$$\frac{dX}{dt} = D(X)\left(G\frac{R}{R+R_0} - d_x\right) \tag{7}$$

$$\frac{dR}{dt} = K - D(d_R)R - D\left(\frac{R}{R+R_0}\right)C^TX \tag{8}$$

where $R_0$ is a constant affinity parameter, $d_x$ is an N $\times$ 1 vector of consumer death rates $d_R$ is an N $\times$ 1 vector of resource depletion rates, G is the growth rate matrix defined in the main text, where $G_{i,\alpha} = r_{i,\alpha}\gamma_{i,\alpha}$, C is the matrix comprised of $r_{i,\alpha}$, where $C_{i,\alpha} = r_{i,\alpha}$, $K$ is the vector of resource supply rates, and D is used as the standard diag operator for vectors, where $D(X)$ refers to the diagonal matrix with the vector $X$ on the diagonal.

For a given vector of resource supply rates, $K$, the equilibrium abundances of consumers, $X^*$, are given by:

$$X^* = \left(C^T\right)^{-1} D\left(\frac{R^*}{R^*+R_0}\right)^{-1}(K - D(d_R)R^*) \tag{9}$$

provided that the matrix $C^T$ has a well-defined left inverse (or left pseudoinverse), which requires that the number of resources $M$ is greater than or equal to $N$, the number of consumers.

In the special case where $M = N$, or the number of resources in the system is equal to the number of consumers, we can assume that the matrix $G$ has a well-defined left inverse. In this case, using the consumer equations, we have that at equilibrium:

$$\frac{R^*}{R^*+R_0} = G^{-1}d_x \tag{10}$$

and

$$R^* = R_0 \frac{G^{-1}d_x}{\mathbb{1} - G^{-1}d_x} \tag{11}$$

where $\mathbb{1}$ is the $M \times 1$ dimensional vector of all 1's.

Substituting these expressions into equation 9, we obtain:

$$X^* = \left(C^T\right)^{-1}\left(\frac{K}{G^{-1}d_x} - \frac{D(d_R)R_0}{\mathbb{1} - G^{-1}d_x}\right) \tag{12}$$

Then, we consider the response to a small environmental fluctuation in which $K \to K + \epsilon$. In this case, the new equilibrium abundance of consumers will be given by:

$$(X^*)' = \left(C^T\right)^{-1}\left(\frac{K+\epsilon}{G^{-1}d_x} - \frac{D(d_R)R_0}{\mathbb{1} - G^{-1}d_x}\right) \tag{13}$$

$$= X^* + \left(C^T\right)^{-1}\left(\frac{\epsilon}{G^{-1}d_x}\right) \tag{14}$$

Assuming the environmental fluctuations are given by an ensemble, $\rho(K)$, with a covariance matrix $\Sigma_K$, we can then write the expected covariance between consumers $\langle X_i, X_j\rangle$, as:

$$\langle X_i, X_j\rangle = \left(C^T\right)^{-1}\left(D(G^{-1}d_x)^{-1}\Sigma_K D(G^{-1}d_x)^{-1}\right)C^{-1} \tag{15}$$

In the limit of infinitely slow environmental fluctuations, a community is expected to equilibrate at an essentially constant influx rate $K$. Thus this expression, while derived for discrete environmental perturbations, is the expected long-term covariance between consumers in the slow fluctuation limit.

**Relationship between covariance matrix and overlap matrix**

To relate the overlap matrix $O = GG^T$ to the covariance matrix, we first note that each entry of the matrix $C$ may be written as $C_{i,\alpha} = \frac{1}{\gamma_{i,\alpha}}G_{i,\alpha}$, where $\gamma_{i,\alpha}$ and $G_{i,\alpha}$ are given as above. If we assume that the yields take a constant value across all species and resources, or that $\gamma_{i,\alpha} = c\ \forall i, \alpha$ for some constant $c$, then the matrix $C$ may be written as $C = \frac{1}{c}G$.

With this, and defining the matrix $Z$ as:

$$Z = \left(D(G^{-1}d_x)^{-1}\Sigma_K D(G^{-1}d_x)^{-1}\right)^{-1}$$

We may then rewrite Eq. 15 as:

$$\langle X_i, X_j \rangle = c^2 (GZG^T)^{-1} \tag{16}$$

While the elements of the matrix $Z$ will in general depend on the exact parameters of the system, as well as the covariance matrix of environmental fluctuations $\Sigma_K$, it is nonetheless clear that the expected long-term covariance between consumers is defined by a scaled version of the matrix $GG^T$.

We consider a simplified scenario in which $R^*_\alpha$, $R_0$ are identical for all resources $R_\alpha$, and the matrix $\Sigma_K$ is diagonal, implying that the covariance between any two influx rate fluctuations, $K_i$ and $K_j$, is 0 for $i \neq j$.

In this case, the matrix $Z = dI$, a constant diagonal matrix, and Eq. 15 may be further simplified as:

$$\langle X_i, X_j \rangle = \frac{c^2}{d} (GG^T)^{-1} \tag{17}$$

If the assumptions of this simplified scenario hold, we expect $GG^T$ and $(GG^T)^{-1}$ to be good approximations for consumer covariances across short and long-timescale fluctuations, respectively. However, deviations from this simplified scenario, such as variance in the values of $R^*_\alpha$ for a given system, will mean that $GG^T$ is likely to become an increasingly poor approximation of the covariance between consumers across all timescales.

#### Cross-feeding interactions produce positive correlations on long timescales

Here, we consider cross-feeding interactions (*63*) in addition to resource competition interactions. Such interactions are common (*1, 41, 64, 65*), and occur when one strain consumes a resource and subsequently excretes a different resource that can be consumed by another strain, such as in overflow metabolism (*30*). We model these interactions here as follows.

$$\begin{aligned}\frac{dx_i}{dt} &= x_i\left(\sum_{\alpha=1}^{M} r_{i,\alpha}\gamma_{i,\alpha}\frac{R_\alpha}{R_\alpha+R_0} - d_x\right)\\ \frac{dR_\alpha}{dt} &= K_\alpha(t) + \sum_{i=1}^{N}\left[-r_{i,\alpha}\frac{R_\alpha}{R_\alpha+R_0}x_i + \sum_{\beta=1}^{M} r_{i,\beta}\frac{R_\beta}{R_\beta+R_0}x_i S_{i,\beta}T_{\alpha,\beta}\right] - R_\alpha d_R\end{aligned} \tag{18}$$

where $S_{i,\beta}$ are the excretion coefficients encoding whether byproducts of resource $\beta$ are excreted by strain $i$, and $T_{\alpha,\beta}$ are the transformation coefficients encoding the fraction of resource $\beta$ that is converted to resource $\alpha$.

We can summarize the cross-feeding interactions between strains by defining two matrices. The first, which we term the cross-feeding matrix $F$, encodes the resources excreted by each strain. In particular,

$$F = ST^T \tag{19}$$

where $S$ and $T$ are defined above, and $F_{i,\alpha}$ quantifies the resource $\alpha$ excreted by strain $i$. Next, we define a cross-feeding flux matrix $J$,

$$J = FC^T \tag{20}$$

where C is the matrix comprised of $r_{i,\alpha}$, where $C_{i,\alpha} = r_{i,\alpha}$. By this definition, $J_{i,j}$ has units of resource/biomass/time and quantifies the flux of resources excreted by strain $i$ and consumed by strain $j$ per unit biomass.

To test the effect of cross-feeding on community response to environmental fluctuations, we numerically integrate Eq. 18 with the sinusoidal resource influx rates of Eq. 6. Our cross-feeding structure simulates trophic cross-feeding, where the orange guild consumes primary resources and supplies them to the blue guild. In particular, each strain in the orange guild excretes resources 41-80 after consuming resources 1-40 ($S_{i\leq 10,\beta\leq 40} = 1$ and $T_{40<\alpha\leq 80,\beta\leq 40} = 1$; $F$ matrix in Fig. S5C). This results in cross-feeding interactions where the orange guild supplies resources to the blue guild

(Fig. S5D). We also set $K_{40<\alpha\leq 80}(t) = 0$, so that the only resources available to the blue guild are supplied by the orange guild. In this trophic cross-feeding case, we see intra-guild cohesion on short timescales, but not on long timescales (correlated dynamics, Fig. S5E). This matches the non-cross-feeding case qualitatively. *Inter*-guild dynamics, however, are correlated across all timescales due to the facilitation of blue guild growth by the orange guild, with these values decreasing to a small but positive correlations at long timescales (Fig. S5E). Notably, this is distinct from the resource competition case, where we do not observe such inter-guild correlations (Fig. 2).

Overall, these results show that intra-guild cohesion is robust to qualitatively different inter-guild interaction mechanisms. Furthermore, the correlations between strains interacting via cross-feeding appear to be correlated on all timescales. This is qualitively distinct from strains interacting via resource competition, which are correlated on short timescales and anti-correlated on long timescales. This observation suggests that inference of interaction type may be made by studying correlation functions between strains subject to differing environmental fluctuation rates.

### Supplementary Figures

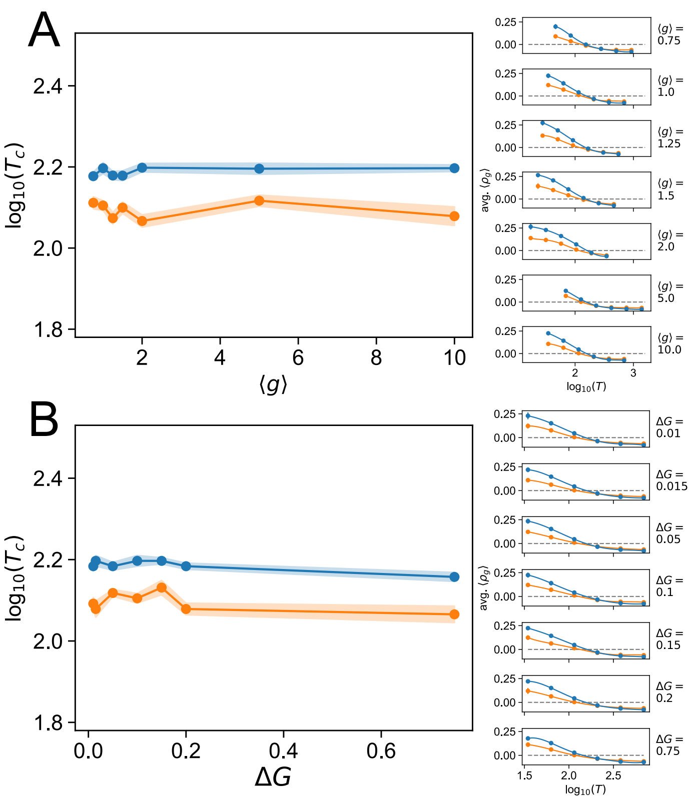


Figure S1: **Growth rate does not set timescale.** **(A)** Average growth rate $\langle g \rangle$ does not set $T_c$ . The left panel shows the logarithm of the crossing timescale $T_c$, plotted as a function of $\langle g \rangle$. $\langle g \rangle$ is varied by varying $\langle r \rangle$, setting $\Delta G = 0.1$ and all other parameter values to the default values (Methods). The shaded region shows uncertainty, calculated via resampling simulations with replacement (10 simulations, Methods). The right panels show the intra-guild correlation $\langle \rho_g \rangle$, averaged across simulations, as a function of the log of the environmental fluctuation timescale $T$ for different values of $\langle g \rangle$. Errorbars indicate standard deviation over simulations. **(B)** Variance in the growth rate matrix $\Delta G$ does not set $T_c$. The left panel shows $\log(T_c)$, plotted as a function of $\Delta G$. Average growth rate $\langle g \rangle$ is set to 1 by changing average uptake rate $\langle r \rangle$ to 5 and keeping yields $\gamma_{i,\alpha}$ at the default value of 0.2, so $\Delta G$ can be interpreted as a fraction of growth rate (Methods). The shaded region shows uncertainty, calculated via resampling simulations with replacement (10 simulations, Methods). The right panels show the average $\langle \rho_g \rangle$ as a function of $T$ for different values of $\Delta G$. Errorbars indicate standard deviation over simulations.

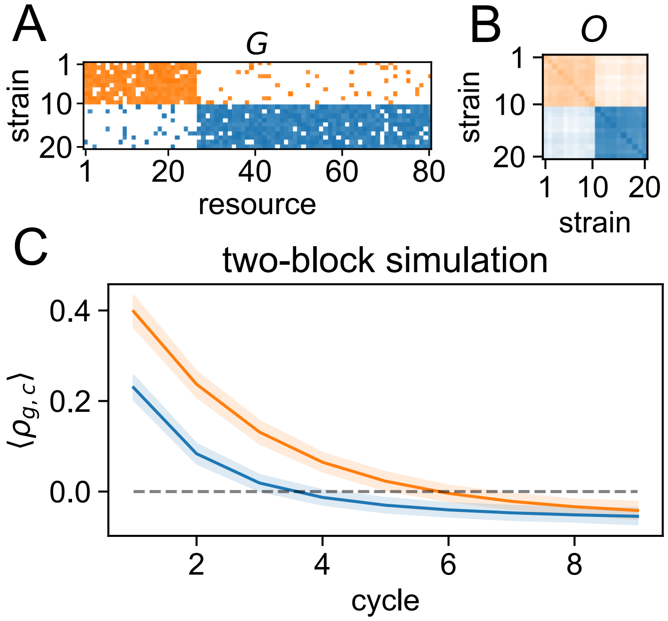


Figure S2: **Dynamic dependence of guild cohesion in batch growth conditions.** Experimental batch culture growth is simulated in a two-block community by numerical integration of Eq. 4 with default batch simulation parameters (Methods). Parameters differ from the simulation of the experiment (Fig. 4E), but reproduce qualitatively the behavior seen in the experimental simulation and experiment. **(A,B)** $G$ and $O$ matrices are generated and shown as described in Fig. 2A and B, except that private resources are omitted. **(C)** Timeseries of the simulated average intra-guild correlation at each cycle across environments. To generate environments, each initial resource value $K_{B,\alpha}$ was set to either $0$ or the default value of $1$, each with probability $0.5$ (Methods). Variance is calculated by resampling with replacement (Methods) and shown by the shaded region. Orange and blue correspond to the orange and blue guilds, respectively (panels A and B).

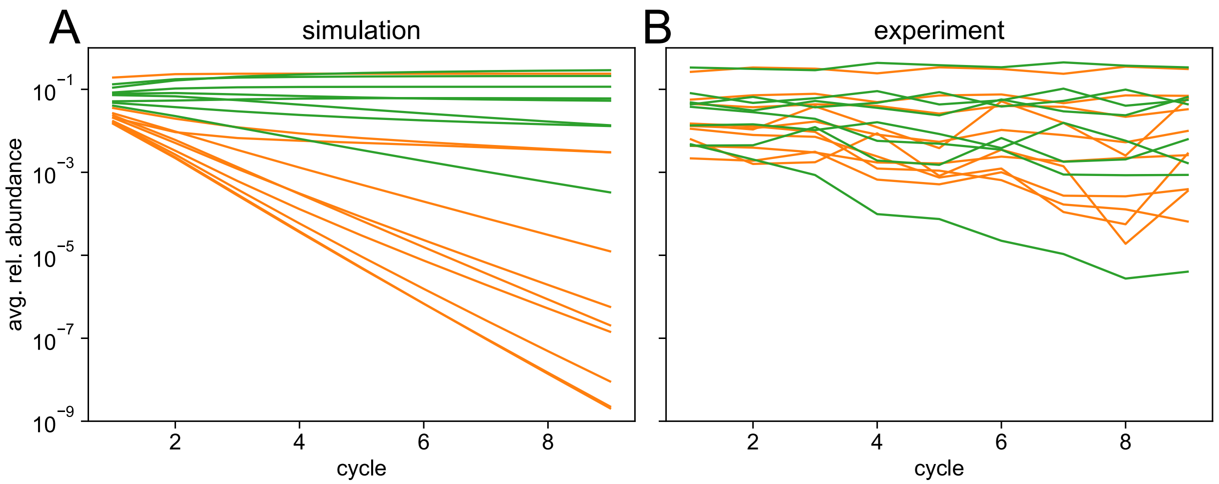


Figure S3: **Average strain relative abundance by guild. (A,B)** Average relative abundance across environments for each synthetic community strain in the green and orange guilds shown in Fig. 4 in simulation (panel A, corresponding to panel D in Fig. 4) and experiment (panel B, corresponding to panel E in Fig. 4). The simulation uses the experimental growth rate matrix $G$, along with the experimental environments (Methods).

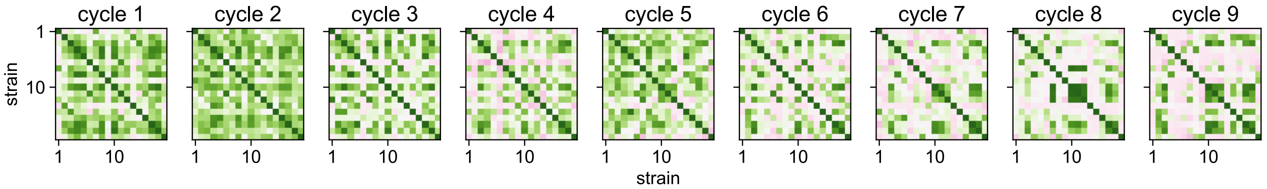


Figure S4: **Experimental correlation coefficients** Strain-strain correlation matrix $\rho_{i,j,c}$ during the experiment shown in Fig. 4. Correlations are calculated based on absolute abundances inferred at the end of each cycle from sequencing with an experimental spike-in (Methods).

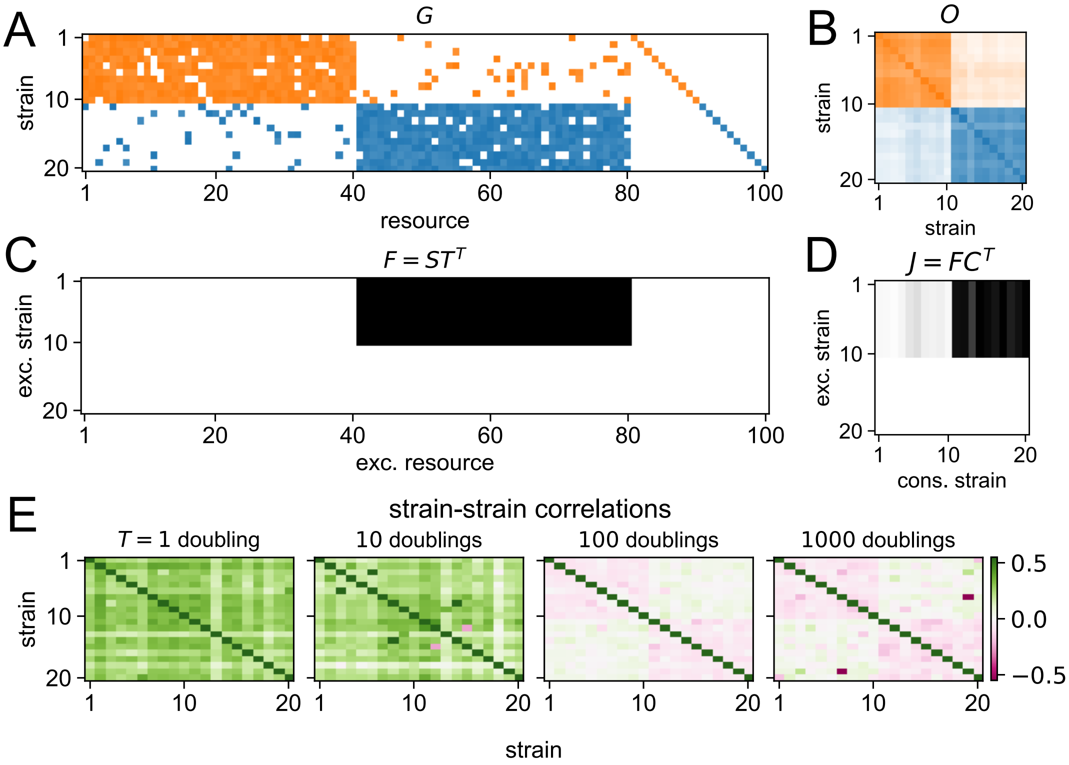


Figure S5: **Cross-feeding gives rise to persistent positive correlations between cross-feeding guilds. (A)** $G$ matrix for cross-feeding simulation, generated and displayed as in Fig. 2A (Eq. 1). **(B)** $O$ matrix for cross-feeding simulation, generated and displayed as in Fig. 2B (Eq. 1). **(C)** Cross-feeding matrix $F = ST^T$. Resources 1-40 are secreted by strains 1-10 and transformed into resources 41-60. The colormap varies linearly from 0 to $\max(F)$. **(D)** Cross feeding flux matrix $J = FC^T$. Resources excreted by strains 1-10 are primarily consumed by strains 11-20. The colormap varies linearly from 0 to $\max(J)$. **(E)** Pearson's correlation coefficient for the abundance dynamics of each strain pair is shown at each fluctuation timescale, as in Fig. 2D.
